# Unexpected false feelings of familiarity about faces are associated with increased pupil dilations

**DOI:** 10.1101/2021.02.22.432360

**Authors:** Kirsten Ziman, Jeremy R. Manning

## Abstract

Our subjective sense that something we encounter is familiar to us is reflected by changes in pupil size. Although pupil dilation effects of familiarity have been well documented, familiarity is not the only, or even the strongest, contributing factor to pupil dilation. Changes in pupil dilation also reflect changes in brightness, affective or emotional responses, hormonal release, expected value or utility, and surprise, among others. Because many factors can affect pupil dilation, important questions remain about how pupil dilation changes reflect high-order cognitive processes, like attention and memory. For example, because surprise and familiarity are often difficult to fully distinguish (since new experiences can be surprising or unexpected), it can be difficult to tease apart pupil dilation effects of surprise versus familiarity. To better understand the effects of surprise and familiarity on pupil dilation, we examined pupil responses during a recognition memory task involving photographs of faces and outdoor scenes. When participants rated novel face images as “familiar,” we observed a robust pupil dilation response.

## Introduction

Imagine that you are observing a crowd of people when you suddenly and unexpectedly notice a childhood friend, whom you haven’t seen in many years, milling amongst the group. You call out and wave, walking towards them. However, when you are able to get a better look, you realize that it isn’t your friend at all–it’s a stranger that you’ve never met before. Awkwardly, you withdraw your hand and pretend to melt back into the scenery.

Research on false memory has shown that people often “fill in” perceived gaps in their recall by building on the scaffolding of their prior knowledge of the current context or situation (Deese, 1959; Roediger & McDermott, 1995; Gallo, 2006; Loftus, 1997). In essence, we pattern complete missing information based on our expectations. But what leads us to mistakenly identify an *unexpected* novel face, place, object, experience, or situation as familiar? We hypothesized that some new insights might come from an unexpected data source: pupil dilations.

Our pupils constrict when we move from a dark setting into a bright one, and dilate when we move from bright to dark. This serves to protect our retina’s photoreceptors in the presense of excessive light energy, and to increase the available light energy when it is more limited. However, this involuntary response is not solely related to the physical intensity or energy of the light shining on the retina. For example, similar pupil constrictions and dilations may also be observed in response to *perceived* brightness or darkness (e.g., in brightness illusions), suggesting that pupillary responses are in part driven by subjective experiences (Laeng & Endestad, 2012). Although brightness is perhaps the strongest driver of the pupillary response, a growing body of work has shown that pupil dilation also tracks with a wide variety of higher-order cognitive processes. For example, pupil dilations also reflect changes in affect and emotion (Oliva & Anikin, 2018; Siegle et al., 2003), attention to high-level information (O. E. Kang et al., 2014), the focus of high-level attention (O. Kang & Wheatley, 2015), synchronization between individuals engaged in conversation (O. Kang & Wheatley, 2017), hormonal release (McCorry, 2007), expected value or expected utility (Slooten et al., 2018), surprise (Preuschoff et al., 2011), and familiarity (Võ et al., 2007; Gardner et al., 1974, 1975).

When pupillary responses reflect high-order cognitive processes, it can be difficult to specifically identify the underlying causes of those responses, in part because many of these high-order processes are inter-related or otherwise inter-dependent. For example, when we encounter something unfamiliar, it can be surprising; evoke a sense of curiosity, fear, joy, or another affective response; cause us to evaluate its expected utility; and so on. Therefore, even well-studied and relatively stable pupillary response effects, such as the finding that our pupils dilate in response to familiar stimuli or experiences (Gardner et al., 1974; Võ et al., 2007; Heaver & Hutton, 2011; Goldinger & Papesh, 2012; Papesh et al., 2012; Naber et al., 2013; Kafkas & Montaldi, 2015; Mill et al., 2016; Jacoby, 1991; Mandler, 1980; Godden & Baddeley, 1975; Vilberg & Rugg, 2008; Yonelinas, 2002) can be difficult to interpret. The recognition memory processes that lead us to feel a sense of familiarity depend in turn on myriad factors and processes that are also associated with changes in pupil dilation (Faber, 2017; Beukema et al., 2019; Zekveld et al., 2018; Kahneman & Beatty, 1966; Kahneman et al., 1967; Ahern & Beatty, 1981; Fiedler & Glöckner, 2012; Einhäuser, 2017).

Here we sought to tease apart the pupillary responses associated with the *feeling* of familiarity from those related to the recognition memory processes that enable us to recognize when a stimulus or experience is *truly* familiar. We designed two conditions of an eyetracking experiment that first asked participants to attend to a series of locations and stimulus features while unattended stimuli and features also appeared on the screen. The two conditions differed in whether the attention cues were consistent across a series of presentations (*Sustained Attention*) or whether they varied randomly with each stimulus presentation (*Variable Attention*). In both conditions, we then asked participants to perform a recognition memory task whereby they rated the “familiarity” of attended, unattended, and novel stimuli. We examined pupillary responses as participants attended different stimuli and as they later made their familiarity judgements. In addition to replicating several previously reported attention-related pupillary response patterns, we also report pupillary responses to *novel* stimuli (i.e., that participants had not seen before) that they nonetheless identified as familiar.

## Materials and methods

We sought to determine if items that feel familiar elicit unique pupil dilation responses, even if they are not truly stored in memory. To answer this question, we leveraged our previously published data from an experiment designed to test the effects of attention on memory. The full dataset may be downloaded [here](https://github.com/ContextLab/attention-memory-task), and the specific experimental groups and conditions we analyzed in the present manuscript may be downloaded [here](https://github.com/ContextLab/attention-memory-task/tree/updates/sustained_attention_experiment/data/group2) and [here](https://github.com/ContextLab/attention-memory-task/tree/updates/variable_attention_experiment/data/group2). All of the analysis code used in our manuscript may be downloaded [here](https://github.com/ContextLab/pupil-memory-analysis).

### Experiment design

The experiment comprised a series of presentation blocks and memory blocks. Throughout the presentation and memory blocks, pupillometry data were collected using an Eyetribe eye-tracking system (Eye Tribe, The EyeTribe, Copenhagen, Denmark). Full experimental and methodological details may be found in Ziman et al. (2020).

#### Presentation blocks

During presentation blocks, participants viewed a series of composite image pairs (one on the left and one on the right of the screen) while keeping their gaze pointed towards a centrally located fixation cross. Each composite image comprised an equal blend of a contrast and brightness normalized grayscale image of a face and an outdoor scene. Participants also received a visual attention cue (Fig. 1a) prior to viewing the composite image pairs (Fig. 1b), directing them to attend to face or scene component (*category*) of the left or right image (*location*). The frequency with which the attention cue was changed varied across two experimental conditions: a *Sustained Attention* and a *Variable Attention* condition.

**Figure 1.**
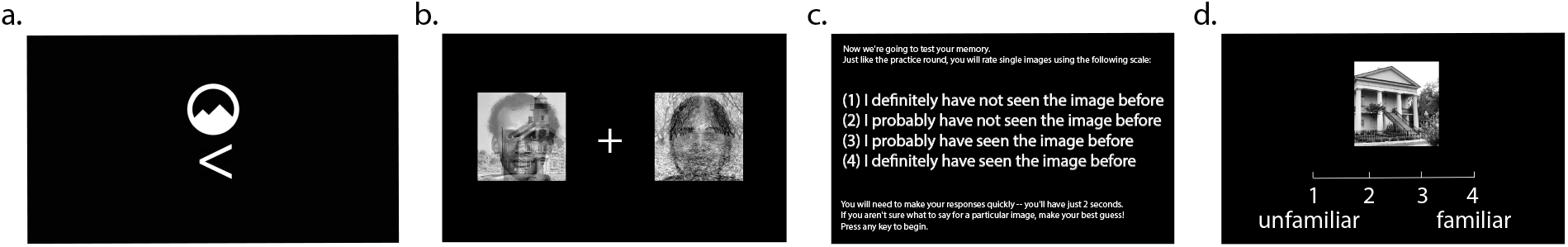
Experimental methods. **a.** During presentation blocks, participants received cues, like the one displayed here, directing their attention to the face or scene component of the left or right composite image. The example cue is directing the participant to attend to the scene component of the left image. **b.** An example composite image pair with a central fixation cross. **c.** Screenshot of instructions shown to participants prior to each memory block. **d.** An example image and familiarity rating response scale displayed during a memory block. Note that the scale of the text and images in all panels have been altered for illustrative purposes.

#### Sustained attention

In the Sustained Attention condition of the experiment (*n* = 30), participants received a single attention cue at the start of each presentation block. In other words, they kept their attention focused on the same image location and category throughout all of the composite image pair presentations. The attention cues were organized across blocks such that location and category were counterbalanced over the course of the experiment.

#### Variable attention

In the Variable Attention condition of the experiment (*n* = 23), participants received a new attention cue prior to every image pair presentation in the presentation block. In other words, they varied the focus of their attention on an image-by-image basis throughout the duration of the presentation block. The location and category cues within and across blocks were counterbalanced over the course of the experiment.

#### Memory blocks

During memory blocks, participants were instructed (Fig. 1c) to rate how “familiar” each of a series of grayscale images seemed on a scale from 1–4. If participants felt unsure about how to respond, they were explicitly instructed to take their best guess. Each image participants judged (Fig. 1d) was drawn either from the set of grayscale face and scene images that they had studied (as part of a composite image pair) during the prior presentation block (*old* images), or from a separate set of images that the participants had not encountered before (*novel* images). The set of images judged in each memory block comprised half old images and half novel images. In turn, the set of old images comprised an equal mix of presented images that whose locations were versus were not attended, and whose categories were versus were not attended. Across all memory blocks, participants viewed (and rated) a total of 80 novel face images and 80 novel scene images.

### Pupillometry data analysis and preparation

Our eyetracking system continuously sampled participants’ eye gaze positions (mean accuracy: 0.5° visual angle; mean precision: 0.1° visual angle root mean squared error) and pupil diameters at 30 Hz. We excerpted three-second windows that began when each new image or composite pair appeared on the participant’s screen. For presentation trials, this window spanned the full duration that composite images displayed on the screen (3s). For memory trials, this window spanned the duration individual images appeared on the screen (2s) in addition to a fixation period after each image disappeared (1s).

We excluded samples where any of the following criteria held: the diameter for either pupil was measured as zero; the inter-pupillary difference in pupil diameters was greater than 1.5 times the interquartile range (across all trials); the gaze position was outside of the border of the display screen; the horizontal or vertical position was greater than 1.5 times the interquartile range from the average gaze location (across all trials); or the same sample was redundantly recorded. When the average sampling rate (for the remaining samples) dropped below 20 Hz, we removed those trials from our analyses. We estimated the pupil diameter (i.e., the pupil dilation response) at each timepoint by averaging the measured left and right pupil diameters. Finally, we converted these averaged diameters into *z*-scored (standard deviation) units within each participant.

We generated a smooth, regularly sampled, timecourse of the pupil dilation responses to each image by fitting a Piecewise Cubic Hermite Interpolating Polynomial (PCHIP; Fritsch & Carlson, 1980) to the pupil diameters from each trial, and sampling from when the trial began until the moment the last viable pupil dilation measurement in the trial was recorded (rounded down to the nearest of 150 evenly spaced timepoints throughout this interval).

#### Segmenting the pupil response timecourse

The pupil response timecourses we observed during different parts of the experiment were often similar across participants. These timecourses often exhibited an initial dilation just after a new image appeared on the participant’s screen, followed by a constriction, and so on. This suggested that different time intervals (relative to the image onset) might reflect different processes of potential interest. For each type of trial we analyzed (presentation trials, memory trial responses to old images, and memory trial responses to new images), we first computed the average pupil response timecourse across all trials and participants. We next computed the average value, *m*, of the average pupil response timecourse during the time interval of interest. We then segmented the pupil response timecourse into consecutive time bins where the pupil dilations were consistently above or below *m*. This yielded a set of “cut points” for the pupil response timecourse (i.e., mean crossings). Finally, we examined the pupil responses within each of these segments to identify potential differences in pupil dilations as a function of the familiarity ratings that the participants assigned (or would later assign) to different images.

## Results

To explore pupillary responses under different attention and memory conditions, we computed the average pupil dilation response timecourses across trials and participants (see *Pupillometry data analysis and preparation*). We first examined pupillary responses as participants attended composite image pairs while keeping their gaze fixed on a central point (Fig. 1b). We reasoned that these pupillary responses might reflect processes related to controlling the focus of feature-based or location-based (spatial) attention, or related to encoding the images into memory. We observed similar response timecourses across both experimental conditions (Sustained Attention, whereby participants were given the same cue for all composite image pairs within in a block; versus Variable Attention, whereby participants were given a new attention cue prior to viewing each image pair; see Fig. 1a for an example attention cue). Figure 2a displays results averaged across both experimental conditions; Figures S1 and S2 display analogous results broken down by condition. The average pupil dilation timecourse we observed when participants viewed the composite image pairs is displayed in Figure 2a. As summarized in Figure 2, participants’ pupil dilation increased when they attended to images that they later recognized (i.e., rated as familiar during the memory phase of the experiment; blue curve in Panel b). When participants attended to images that they would later fail to recognize, their pupils did not dilate as much (red curve in Panel b). This suggests that participants’ pupils were dilating when they successfully encoded an image from the attended location and category into memory. When we examined pupil responses to unattended images (i.e., images from the unattended location or category) we observed no reliable differences in participants’ pupil response timecourses as a function of the familiarity ratings they assigned to those images during the memory phase of the experiment (Fig. 2c–e). This suggests that the unattended images may not have been encoded into memory as reliably, or that some other mechanism or process that does not track as closely with pupil dilations might govern the encoding of the unattended images.

**Figure 2.**
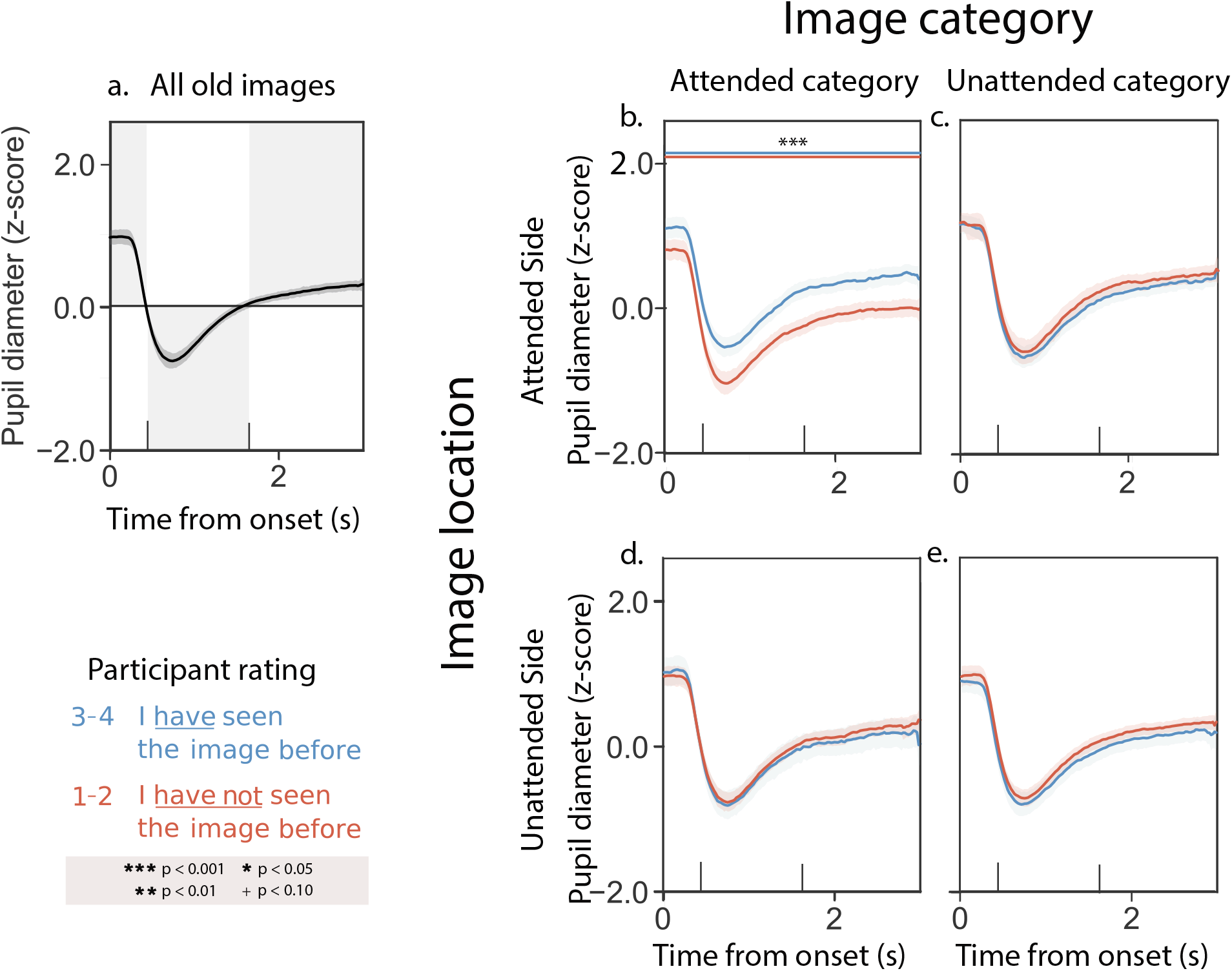
Pupil dilation response timecourses while attended to composite image pairs. **a.** Average pupil dilation timecourse across all trials and experimental conditions. **b.** Pupil dilation timecourses for trials corresponding to attended images that participants later rated as familiar (blue curve; familiarity rating = 3 or 4) versus unfamiliar (red curve; familiarity rating = 1 or 2). **c.** Pupil dilation timecourses for trials corresponding to images on the attended side (but the unattended category) that participants later rated as familiar (blue) or unfamiliar (red). **d.** Pupil dilation timecourses for trials corresponding to images from the attended category (but on the unattended side) that participants later rated as familiar (blue) or unfamiliar (red). **e.** Pupil dilation timecourses for trials corresponding to images on the unattended side, from the unattended category, that participants later rated as familiar (blue) or unfamiliar (red). All panels: error ribbons denote 95% confidence intervals across participants. See Supplemental Figures S1 and S2 for analogous results broken down by experimental condition and numerical familiarity rating. In this figure, and in subsequent figures, the horizontal line pairs denote reliable separation (quantified using two-tailed paired t-tests) between the corresponding curves, during the time intervals covered by the lines. Significance levels are denoted by the symbols shown in the legend.

Next, we examined pupillary responses as participants rated the familiarity of the images that had comprised the composite pairs they had seen during the presentation phase of the experiment. We reasoned that these pupillary responses might reflect processes related to memory retrieval. We observed similar timecourses across both the Sustained Attention and Variable Attention experimental conditions. Figure 3a displays results averaged across both experimental conditions; Figures S3 and S4 display analogous results broken down by condition. The average pupil dilation timecourse we observed when participants viewed previously seen memory cue images is displayed in Figure 3a. As summarized in Figure 3, participants’ pupil dilation increased (numerically) when they recognized previously attended images as familiar (blue curve in Panel a) versus when they failed to recognize previously attended images as familiar (red curve in Panel a). We observed a qualitatively similar increase in pupil dilation when participants rated partially attended images as familiar (blue curves in Panels c and d) versus unfamiliar (red curves in Panels c and d). Although the responses displayed in Panels b–d are all qualitatively similar, only the differences in Panel d crossed our threshold for statistical significance. Finally, we saw no consistent familiarity-dependent changes in pupil responses to unattended images (Panel e).

**Figure 3.**
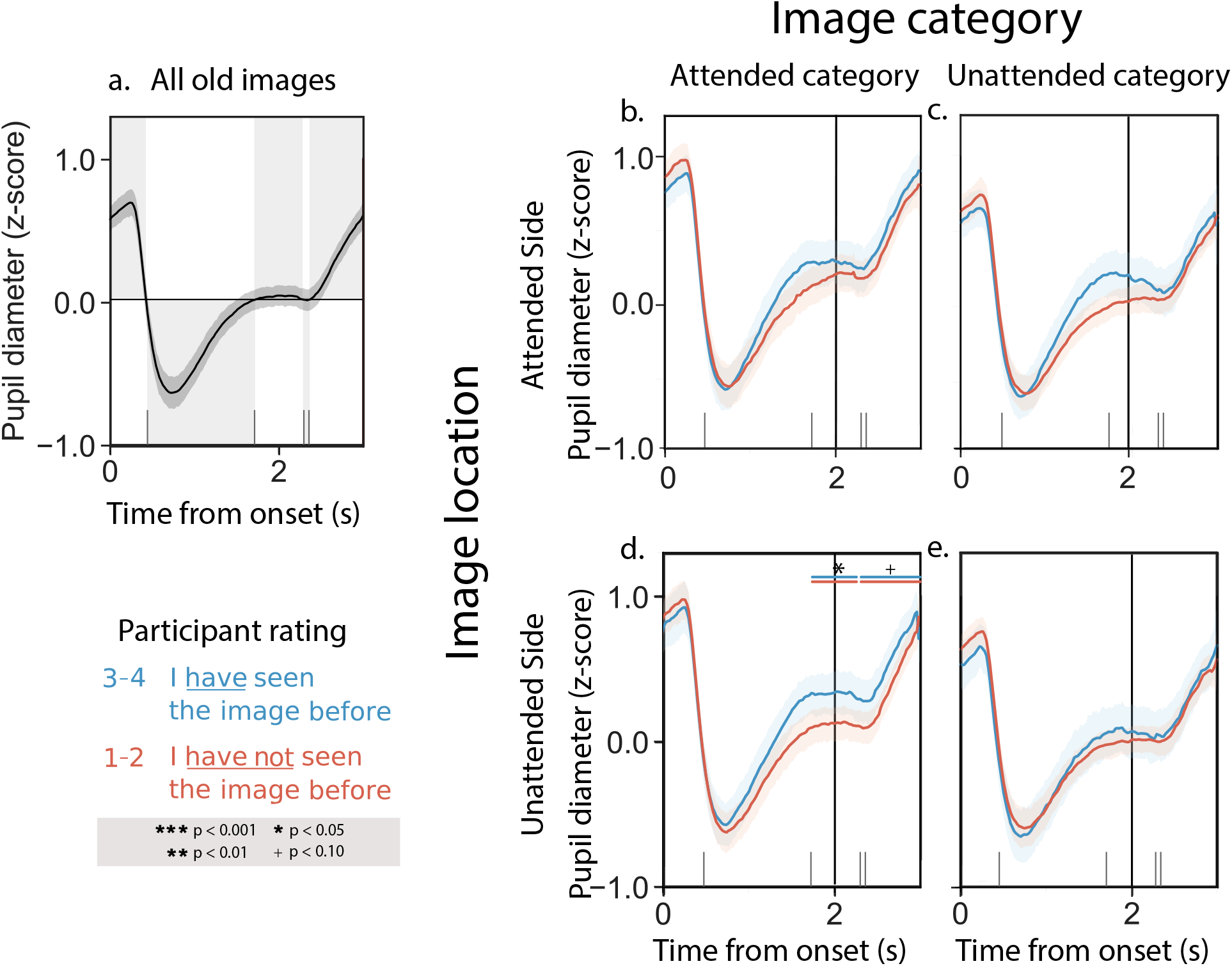
Pupil dilation response timecourses while rating the familiarities of previously studied images. **a.** Average pupil dilation timecourse across all trials and experimental conditions. **b.** Pupil dilation timecourses for trials corresponding to previously attended images that participants rated as familiar (blue curve; familiarity rating = 3 or 4) versus unfamiliar (red curve; familiarity rating = 1 or 2). **c.** Pupil dilation timecourses during trials where participants rated images on the attended side (but the unattended category) as familiar (blue) or unfamiliar (red). **d.** Pupil dilation timecourses during trials where participants rated images from the attended category (but the unattended side) as familiar (blue) or unfamiliar (red). **e.** Pupil dilation timecourses during trials where participants rated unattended images as familiar (blue) or unfamiliar (red). All panels: error ribbons denote 95% confidence intervals across participants. The vertical lines indicate when the images were cleared from the screen. See Supplemental Figures S3 and S4 for analogous results broken down by experimental condition and numerical familiarity rating.

Taken together, the above pattern of results could be consistent with several possible interpretations. One possibility is that participants’ pupils dilate during memory retrieval, analogous to the responses we observed during the presentation phase of the experiment that appeared to track with memory encoding. This seems to be supported by the finding that differences in pupil dilation responses to images that were rated as familiar versus unfamiliar appear to fall off monotonically as a function of how much attention participants were instructed to pay to the corresponding images during the presentation phase of the experiment (e.g., compare Panels b–d with Panel e). In this way, our results thus far potentially agree with findings from myriad studies showing that people’s pupils dilate when they are engaged in remembering or recognizing (Goldinger & Papesh, 2012; El Haj et al., 2019; Rijn et al., 2012; Kucewicz et al., 2018; Naber et al., 2013; Mill et al., 2016). However, an alternative explanation is that pupil dilations might instead reflect the *feeling* of remembering or recognizing as opposed to memory retrieval per se. We hypothesized that participants’ familiarity judgements of novel (never before seen) images might enable us to disentangle these explanations. In particular, if we observed a pupil dilation response during the rare times when participants mistakenly rated novel images as familiar, this would indicate that the pupil response is driven in part by the feeling of familiarity rather than the specific engagement of memory retrieval processes.

When we examined participants’ familiarity ratings of novel stimuli, we noticed several behavioral patterns. In the Sustained Attention condition, participants rated novel images as more familiar if they came from the most recently attended category (familiarity ratings of novel stimuli that matched versus conflicted with the most recent attention cue: *t*(29) = 4.37,*p* < 0.001). In the Variable Attention condition, participants tended to rate novel scene images as more familiar than face images, regardless of the most recent attention cue, although this tendency did not cross our threshold for statistical significance (*t*(22) = 1.24,*p* = 0.23). This suggests that when participants modulate their focus of attention to specific stimuli, the consequences to how they subsequently process images linger beyond the duration that the cues remain relevant. When attention cues were stable (i.e., in the Sustained Attention condition), participants responded in a biased way to novel images that matched the most recent stable cue. However, when the attention cues changed rapidly (i.e., in the Variable Attention condition), participants appeared to “default” to processing scenes and face images slightly differently, regardless of the most recent cued category. This suggests that different image categories may be processed or prioritized (in attention, memory, etc.) differently, independent of the specific experimental task, cues, or instructions. We therefore separated our further analyses of responses to novel stimuli along two dimensions: (1) whether or not the novel stimuli came from the most recently cued category and (2) whether the novel stimuli were scene versus face images.

Participants’ pupillary responses to novel stimuli in the Sustained and Variable Attention conditions were similar. Figure 4 displays results averaged across both experimental conditions, and Figures S5 and S6 display analogous results broken down by condition. The average pupil dilation timecourse we observed when participants viewed novel memory cue images is displayed in Figure 4a. Unlike their responses to composite images during the presentation phase of the experiment, or to memory cues for previously seen images during the memory phase of the experiment, participants’ pupillary responses to novel memory cues did not vary reliably as a function of the most recent attention cue (e.g., compare Fig. 4b versus d and c versus e). However, we did observe differences in participants’ pupillary responses as a function of the category (scene versus face) of the novel memory cues. When participants viewed novel scene images, their pupillary responses showed no reliable differences as a function of the familiarity ratings participants assigned to those images (Fig. 4b and d). However, when participants viewed novel face images, their pupils dilated more when they rated the novel images as familiar (Fig. 4c and e).

**Figure 4.**
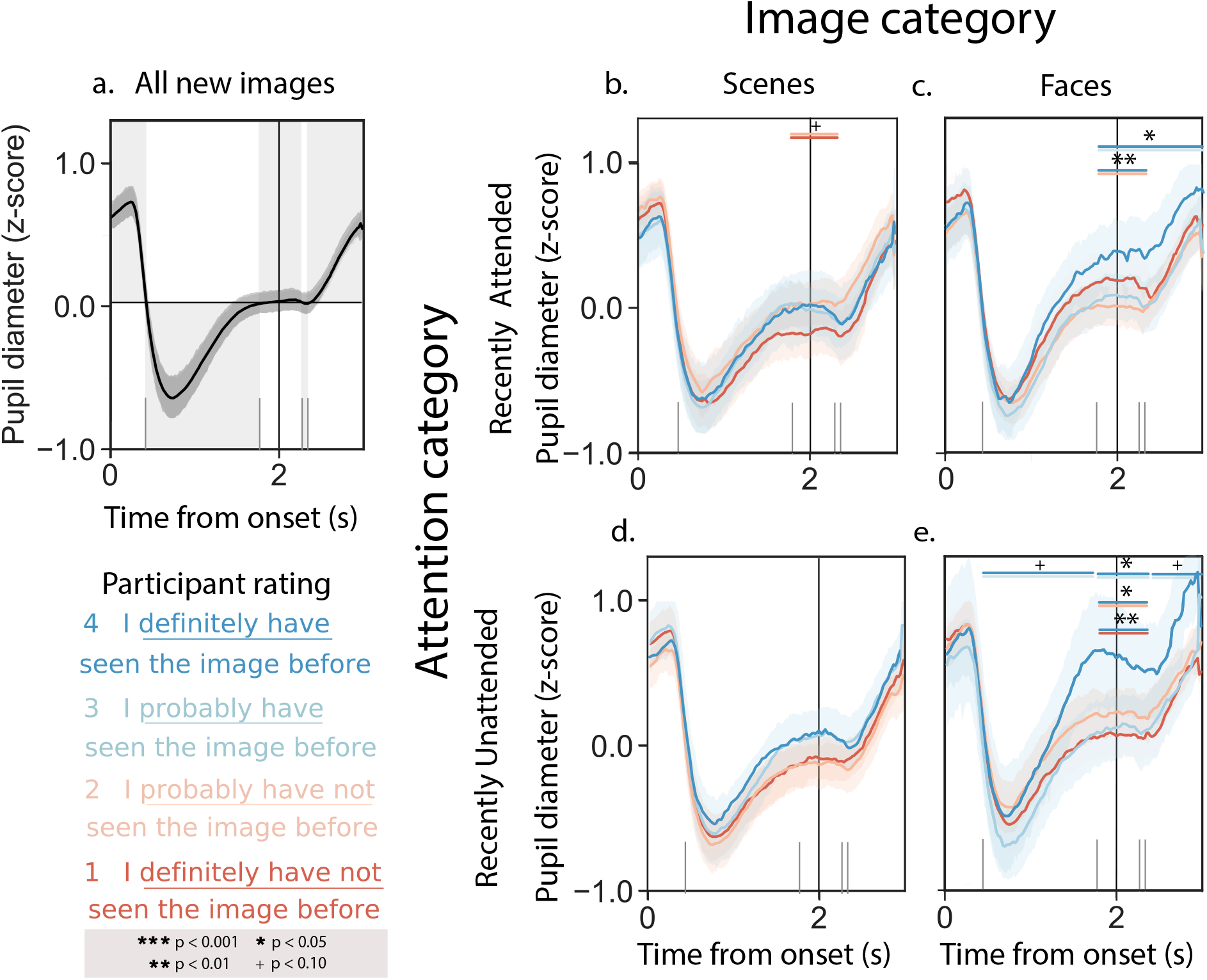
Pupil dilation response timecourses while rating the familiarities of novel images. **a.** Average pupil dilation timecourse across all trials and experimental conditions. **b.** Pupil dilation timecourses (split by familiarity rating) for trials corresponding to novel scene images, when the most recent attention cue was also to a scene image. **c.** Pupil dilation timecourses (split by familiarity rating) for trials corresponding to novel face images, when the most recent attention cue was also to a face image. **d.** Pupil dilation timecourses (split by familiarity rating) for trials corresponding to novel scene images, when the most recent attention cue was to a face image. **e.** Pupil dilation timecourses (split by familiarity rating) for trials corresponding to novel face images, when the most recent attention cue was to a scene image. All panels: error ribbons denote 95% confidence intervals across participants. The vertical lines indicate when the images were cleared from the screen. See Supplemental Figures S5 and S6 for analogous results broken down by experimental condition.

## Discussion

We examined pupillary responses as participants modulated their attention and rated the familiarity of previously seen and novel images. Whereas familiarity and retrieval are often conflated (e.g., when we recognize something we experienced in the past), examining pupillary responses to *novel* stimuli enabled us to disambiguate familiarity and retrieval. When participants rated novel faces as familiar, we observed a pupil dilation response that was qualitatively similar to the pupil dilation response we observed when participants correctly recognized previously encountered stimuli as familiar. However, the pupil dilation response to novel stimuli could not be explained by pure memory retrieval (since there were no prior memories about the stimuli to retrieve), nor could it be explained by response bias (since participants were biased to rate *scenes* as slightly more familiar than faces, all else being equal). Taken together, our findings suggest that the pupil dilation responses we observed are due to participants’ *feelings* of familiarity. Further, this effect seemed specific to participants’ responses to images of faces, in that we did not observe a familiarity-associated pupillary response when participants rated novel scene images.

We note several potential limitations of our study. The most substantial limitation we see is that we cannot entirely rule out that novel stimuli might trigger some sort of partial memory retrieval process. For example, a given novel image might *remind* a participant of other images they had encountered earlier on in the experiment. This could be driven by visual similarity, semantic similarity, or even associations drawn from the participants’ prior experiences. This potential confound means that we cannot completely rule out that the pupil dilations we observed when participants rated novel faces as familiar might be driven in part by memory retrieval processes. However, any such process would need to be category selective, since we did not observe a pupil dilation response to novel scene images (regardless of their familiarity ratings). A second potential limitation of our study is that we cannot distinguish whether the pupil dilation response to familiar-seeming novel faces is specific to faces in particular, or whether it is instead category selective. To distinguish these possible explanations, one would need to collect additional data using images selected from a broader range of categories.

Our study contributes to a growing literature on pupillary responses in a wide range of cognitive tasks, particularly those aimed at studying processes underlying attention and memory (Korn & Bach, 2016). Prior work has also shown that our pupils dilate when we identify a target amidst a distracting background (Wang et al., 2020; Martin & Johnson, 2015), or when we detect an unexpected visual change (Kloosterman et al., 2015). Pupillary responses also track with internal belief states (Colizoli et al., 2018) and pre-conscious processes (Laeng et al., 2012). These findings help to contextualize our finding that participants’ pupils dilated when they rated novel faces as familiar, even though they displayed an overall bias to rate novel faces as unfamiliar. The variety of cognitive phenomena that have been tied to pupillary responses also highlight the richness and complexity underlying the pupillary response. That a scalar value (pupil diameter) at a given moment incorporates such complexity also illustrates how difficult it can be to tease apart the many contributing factors. This also limits our ability to fully interpret pupillometry data (e.g., compared with pure behavioral data, or some other biophysiological measurements under appropriate conditions).

The false feelings of familiarity our participants occasionally exhibited are also informed by a large literature on false memories (Deese, 1959; Roediger & McDermott, 1995; Gallo, 2006; Loftus, 1997). Faces can be an especially interesting stimulus in these experiments given their special relevance and importance to everyday human life. Prior work on recognition memory for face images has shown that feelings of familiarity versus true memory retrieval-based recognition can be dissociated (e.g., by inverting the images Megreya & Burton, 2007), suggesting that these processes may be supported by different mechanisms. Other work has shown that familiarity can also be influenced by visual properties of the faces themselves (e.g., their visual distinctiveness Lewis, 2010). Taken together, this work suggests that the feeling that something is familiar can be at least partially dissociated from remembering that something has been encountered before.

## Acknowledgements

We acknowledge useful discussions with the EPSCoR Attention Consortium, Megan deBettencourt, Caroline Lee, Paxton Fitzpatrick, Sharif Saleki, and Michael Ziman. This work was supported in part by NSF EPSCoR award number 1632738. the views reflected in this manuscript do not necessarily reflect the views of our supporting organizations.

## Notes

https://github.com/ContextLab/pupil-memory-analysis

### Competing Interest Statement

The authors have declared no competing interest.

