## Supporting materials for "Unexpected false feelings of familiarity about faces are associated with increased pupil dilations"

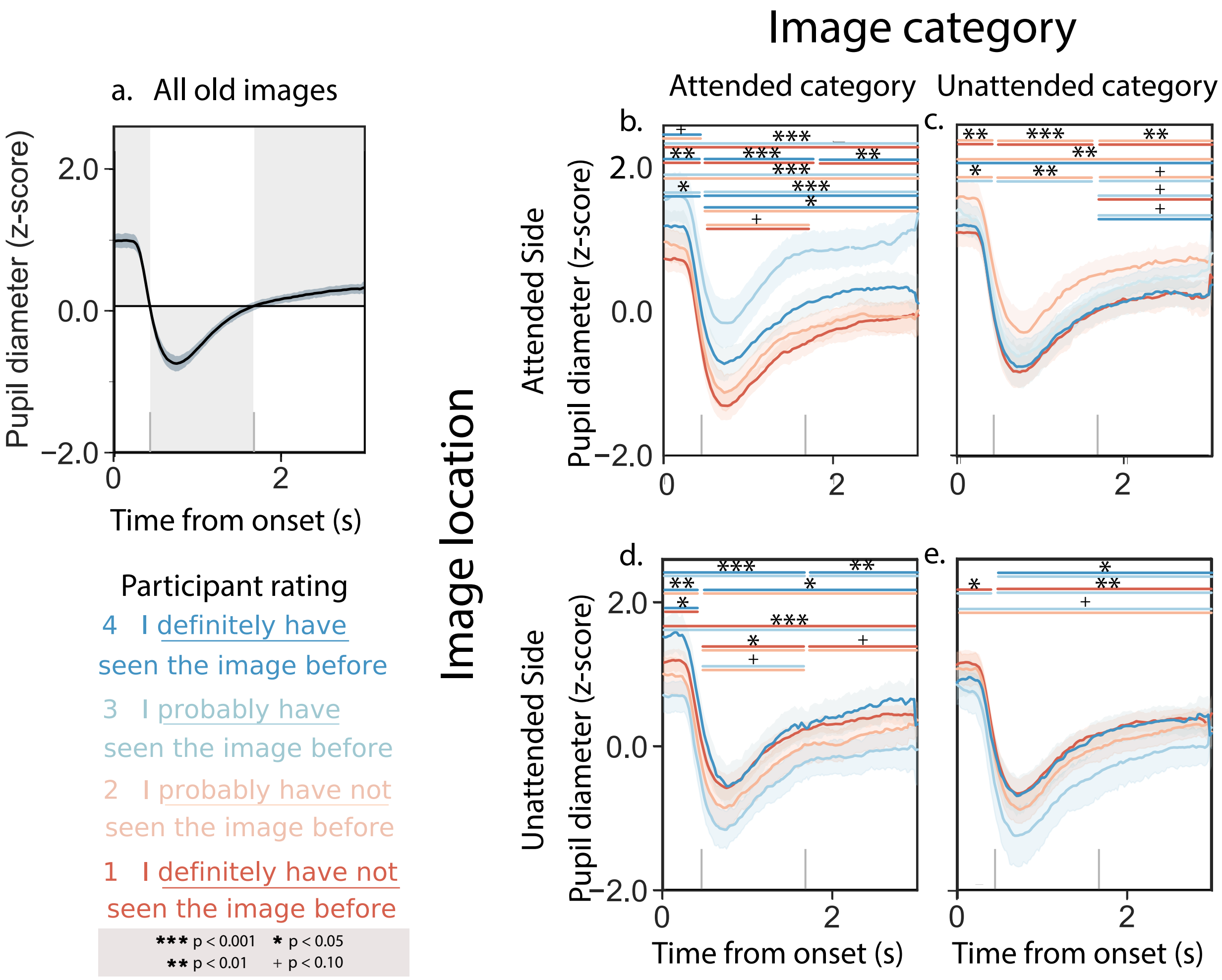


*Figure S1.* **Sustained Attention condition: Pupil dilation response timecourses while attending to composite image pairs. a.** Average pupil dilation timecourse across all trials in both experiments. **b.** Pupil dilation timecourses (split by familiarity rating) for trials corresponding to attended images. **c.** Pupil dilation timecourses (split by familiarity rating) for trials corresponding to images on the attended side. **d.** Pupil dilation timecourses (split by familiarity rating) for trials corresponding to images from the attended category (but on the unattended side). **e.** Pupil dilation timecourses (split by familiarity rating) for trials corresponding to images on the unattended side, from the unattended category. All panels: error ribbons denote 95% confidence intervals across participants. See Figure 2 in the main text for results aggregated across experimental conditions.

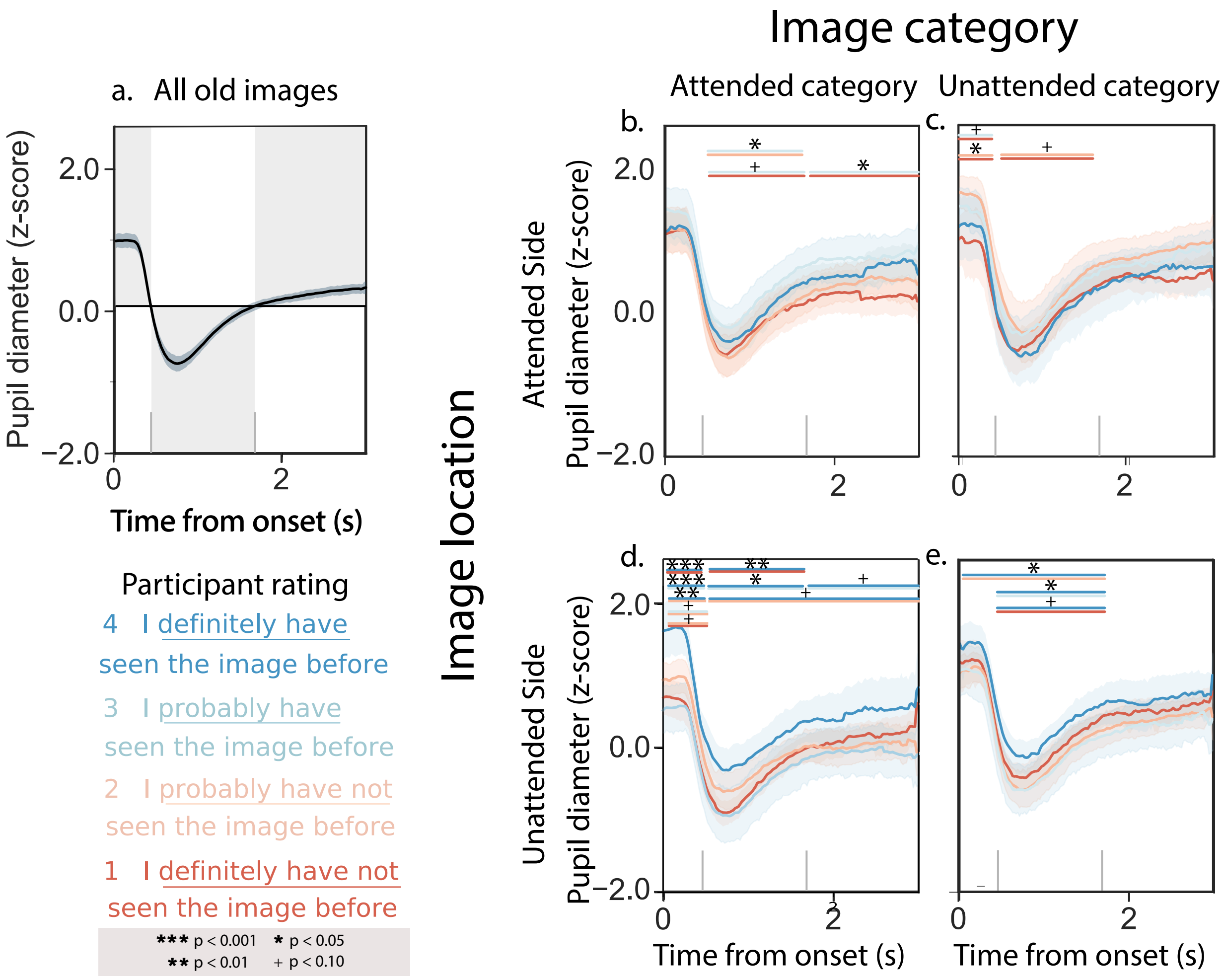


*Figure S2.* **Variable Attention condition: Pupil dilation response timecourses while attending to composite image pairs. a.** Average pupil dilation timecourse across all trials in both experiments. **b.** Pupil dilation timecourses (split by familiarity rating) for trials corresponding to attended images. **c.** Pupil dilation timecourses (split by familiarity rating) for trials corresponding to images on the attended side (but the unattended category). **d.** Pupil dilation timecourses (split by familiarity rating) for trials corresponding to images from the attended category (but on the unattended side). **e.** Pupil dilation timecourses (split by familiarity rating) for trials corresponding to images on the unattended side, from the unattended category. All panels: error ribbons denote 95% confidence intervals across participants. See Figure 2 in the main text for results aggregated across experimental conditions.

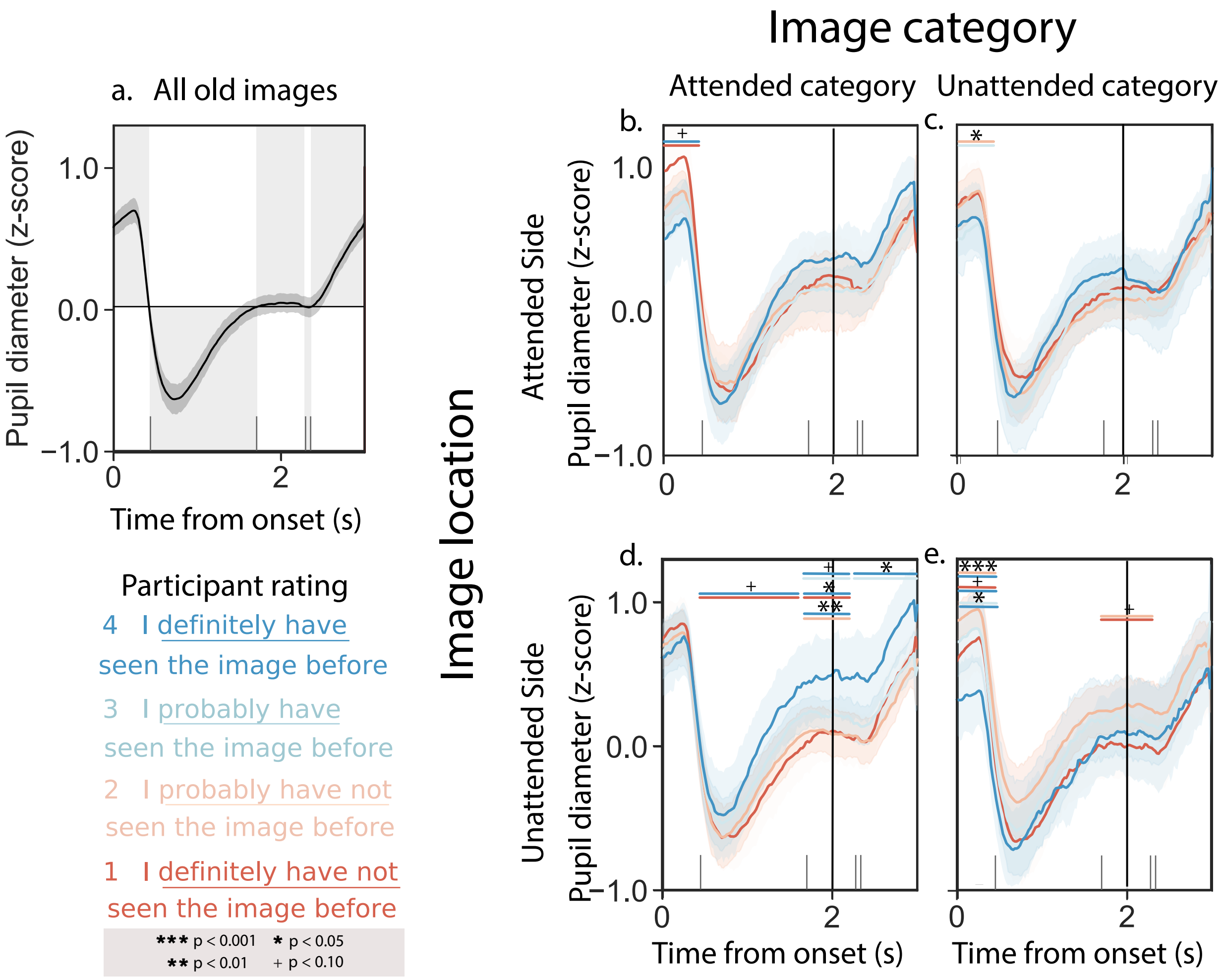


*Figure S3.* **Sustained Attention condition: Pupil dilation response timecourses while rating the familiarities of previously studied images. a.** Average pupil dilation timecourse across all trials in both experiments. **b.** Pupil dilation timecourses (split by familiarity rating) for trials corresponding to previously attended images. **c.** Pupil dilation timecourses (split by familiarity rating) during trials where participants rated images on the attended side. **d.** Pupil dilation timecourses (split by familiarity rating) during trials where participants rated images from the attended category. **e.** Pupil dilation timecourses (split by familiarity rating) during trials where participants rated unattended images. All panels: error ribbons denote 95% confidence intervals across participants. The vertical lines indicate when the images were cleared from the screen. See Figure 3 in the main text for results aggregated across experimental conditions.

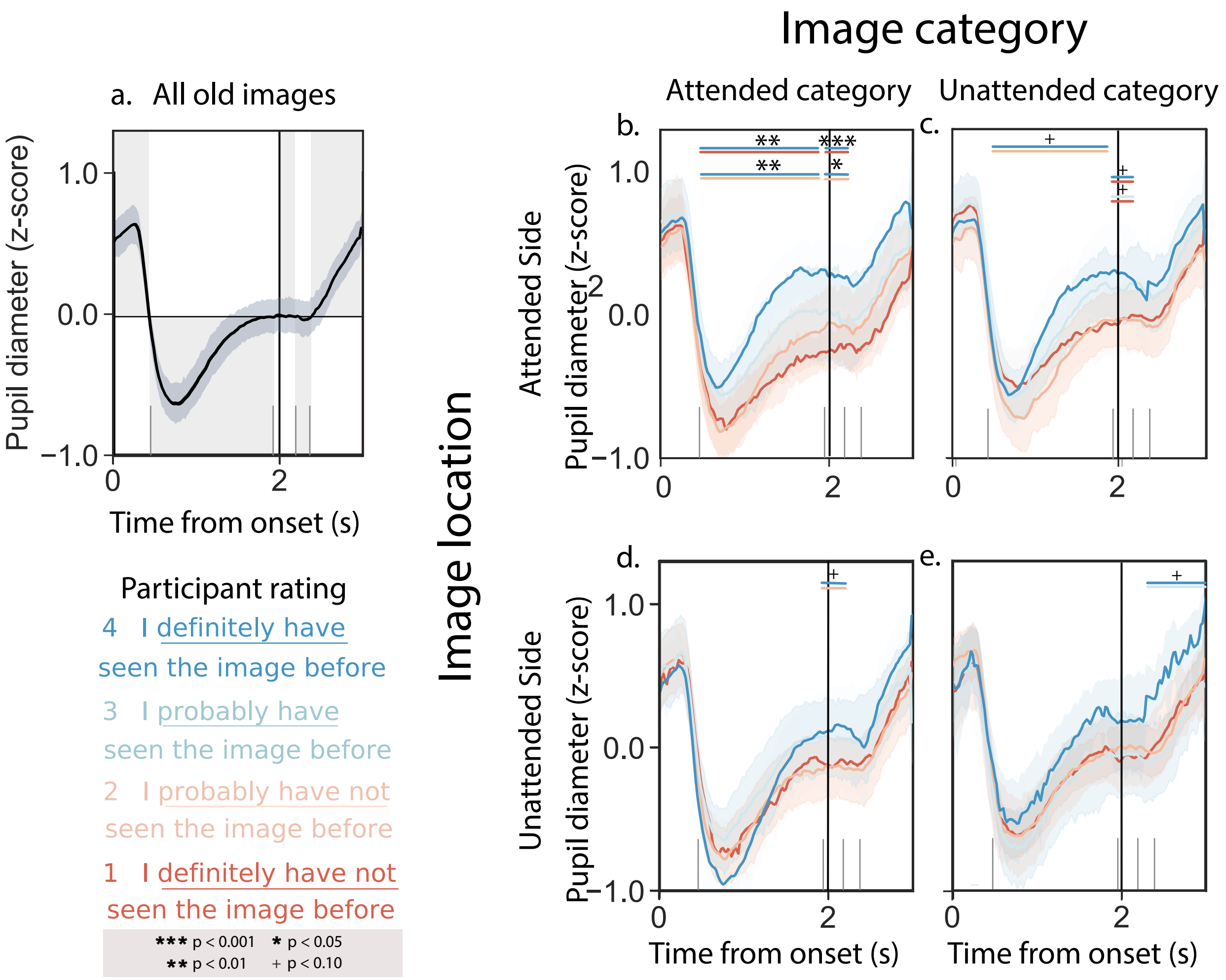


*Figure S4.* **Variable Attention condition: Pupil dilation response timecourses while rating the familiarities of previously studied images.** **a.** Average pupil dilation timecourse across all trials in both experiments. **b.** Pupil dilation timecourses (split by familiarity rating) for trials corresponding to previously attended images. **c.** Pupil dilation timecourses (split by familiarity rating) during trials where participants rated images on the attended side. **d.** Pupil dilation timecourses (split by familiarity rating) during trials where participants rated images from the attended category. **e.** Pupil dilation timecourses (split by familiarity rating) during trials where participants rated unattended images. All panels: error ribbons denote 95% confidence intervals across participants. The vertical lines indicate when the images were cleared from the screen. See Figure 3 in the main text for results aggregated across experimental conditions.

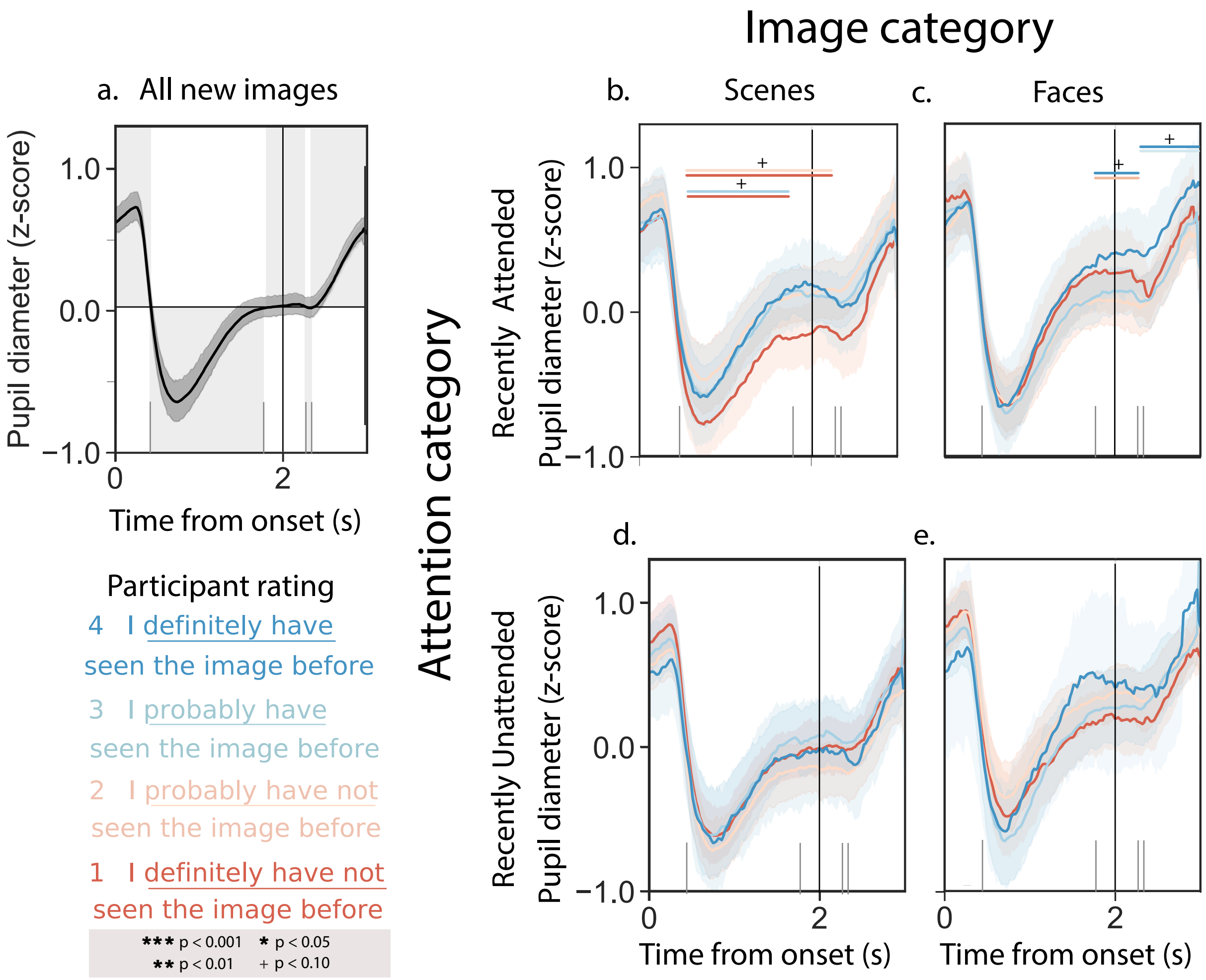


*Figure S5.* **Sustained Attention condition: Pupil dilation response timecourses while rating the familiarities of novel images. a.** Average pupil dilation timecourse across all trials in both experiments. **b.** Pupil dilation timecourses (split by familiarity rating) for trials corresponding to novel scene images, when the most recent attention cue was also to a scene image. **c.** Pupil dilation timecourses (split by familiarity rating) for trials corresponding to novel face images, when the most recent attention cue was also to a face image. **d.** Pupil dilation timecourses (split by familiarity rating) for trials corresponding to novel scene images, when the most recent attention cue was to a face image. **e.** Pupil dilation timecourses (split by familiarity rating) for trials corresponding to novel face images, when the most recent attention cue was to a scene image. All panels: error ribbons denote 95% confidence intervals across participants. The vertical lines indicate when the images were cleared from the screen. See Figure 4 in the main text for results aggregated across experimental conditions.

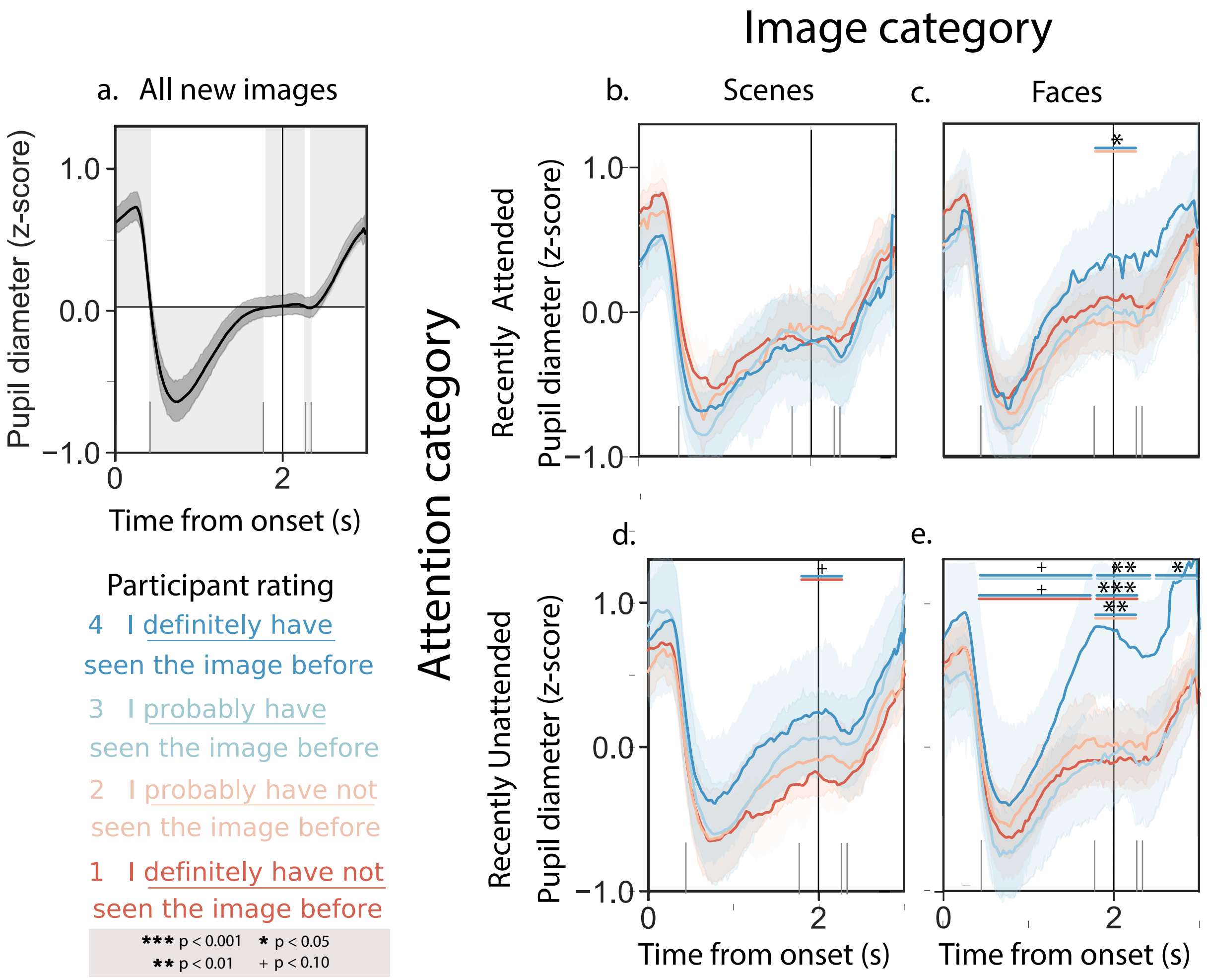


*Figure S6.* **Variable Attention condition: Pupil dilation response timecourses while rating the familiarities of novel images. a.** Average pupil dilation timecourse across all trials in both experiments. **b.** Pupil dilation timecourses (split by familiarity rating) for trials corresponding to novel scene images, when the most recent attention cue was also to a scene image. **c.** Pupil dilation timecourses (split by familiarity rating) for trials corresponding to novel face images, when the most recent attention cue was also to a face image. **d.** Pupil dilation timecourses (split by familiarity rating) for trials corresponding to novel scene images, when the most recent attention cue was to a face image. **e.** Pupil dilation timecourses (split by familiarity rating) for trials corresponding to novel face images, when the most recent attention cue was to a scene image. All panels: error ribbons denote 95% confidence intervals across participants. The vertical lines indicate when the images were cleared from the screen. See Figure 4 in the main text for results aggregated across experimental conditions.
